# FLASH reduces radiation-induced oral mucositis in a mouse model of Fanconi anemia

**DOI:** 10.64898/2026.05.25.727748

**Authors:** Phoebe Loo, Margaret Pan, Man Zhao, Stavros Melemenidis, Dixin Chen, Lucy Whitmore, Sara Richter, Frederick M. Dirbas, Kerriann M. Casey, Edward E. Graves, Michael W. Epperly, Joel Greenberger, Billy W. Loo, Erinn B. Rankin

## Abstract

Patients with Fanconi anemia (FA) are particularly susceptible to developing squamous cell carcinoma of the head and neck due to impaired DNA repair pathways. However, their hypersensitivity to DNA damaging agents can limit effective treatment with standard radiotherapy due to severe side effects and complications. In pre-clinical models, ultra-rapid FLASH radiotherapy (FLASH) reduces radiation-induced toxicity in normal tissues while maintaining similar tumor control compared to conventional dose rate radiotherapy (CONV). Here, we investigated the safety of FLASH for treatment of the head and neck in a mouse model of FA. 129/Sv wild-type (WT) and *Fanca*-deficient (*Fanca*^-/-^) mice received single-dose oral cavity irradiation with electron beam FLASH or CONV to evaluate radiation-induced toxicity in non-tumor-bearing mice. *Fanca* WT and *Fanca*^-/-^ mice were irradiated with 25 and 18 Gy, respectively, of FLASH (190 Gy/sec) or CONV (0.2 Gy/sec), with tongues harvested at 12 hours (hpi) and 10 days (dpi) post-irradiation. At 10 dpi, FLASH-irradiated tongues in both genetic backgrounds demonstrated reduced ulceration at the dorsal tongue surface compared to CONV-irradiated counterparts. Histopathological analysis of the tongue revealed lower mucositis severity scores with decreased epithelial thinning and ulceration in FLASH-irradiated tongues compared to CONV-irradiated ones. Analysis of γ-H2AX foci formation at 12 hpi demonstrated fewer foci in WT mice treated with FLASH compared to CONV, with a similar trend observed in *Fanca*^-/-^ mice. These findings suggest a potential normal tissue-sparing effect with FLASH and hold important clinical implications for the treatment of patients with Fanconi anemia and head and neck cancers.

## Introduction

Ultra-rapid FLASH radiotherapy (FLASH) is emerging as an exciting new treatment paradigm to enhance the therapeutic index of radiotherapy. In preclinical studies, FLASH reduces radiation-induced toxicities in multiple tissues including the lung [1], intestine [2,3], skin [4,5], brain [6,7] and oral cavity [8] compared to conventional radiotherapy (CONV), in the 2-3 orders of magnitude lower delivery rates used clinically, while demonstrating comparable tumor control [1,9]. These findings have generated considerable interest in the potential clinical translation of FLASH as a strategy to expand the therapeutic index of radiotherapy with clinical trials ongoing and in development [10]. However, the mechanisms by which FLASH selectively reduces radiation-induced toxicity to normal tissues remain poorly understood. Moreover, although preclinical studies provide increasing evidence of the advantages of FLASH over CONV, it remains unknown whether these benefits extend to patient populations with intrinsic radiosensitivity.

One such radiosensitive population is patients with Fanconi anemia (FA), an autosomal recessive disorder caused by germline mutations in any of at least 22 FA genes [11,12], which collectively play critical roles in DNA repair [13–15]. FA is characterized by cellular hypersensitivity to DNA crosslinking agents [16] and a markedly increased risk of cancer, with a 50-fold higher incidence of solid tumors and a 500-fold higher incidence of head and neck squamous cell carcinomas (HNSCC) compared to the general population [17–19]. As survival of FA patients has improved through hematopoietic stem cell transplantation, the incidence of malignancies such as HNSCC has become an increasing clinical concern [17]. Since FA patients are highly sensitive to DNA damaging agents, radiotherapy (a key component of standard HNSCC treatment) is often limited by severe normal tissue toxicities including mucositis, dysphagia, and pancytopenia, which can compromise treatment efficacy and overall survival [20–22]. These findings highlight the unmet clinical need for improved treatment strategies to enhance the therapeutic index of radiotherapy for the treatment of solid tumors that develop in FA patients.

The most common FA mutation occurs in the *FANCA* gene, which encodes a protein that functions in the nucleus to coordinate multiple repair processes such as DNA lesion recognition, incision, bypass, and repair, and interacts with other pathways including homologous recombination [12,23–26]. Given its clinical prevalence and milder phenotype relative to other FA mutations [13], *Fanca*-deficient (*Fanca*^-/-^) mouse models provide a clinically relevant model for studying radiosensitivity and strategies to mitigate normal tissue injury during radiotherapy.

Here we investigate FLASH as a strategy to reduce normal tissue toxicities associated with oral cavity irradiation in *Fanca*^-/-^ mice. We found that FLASH reduced oral mucositis in both wild-type (*Fanca*^+/+^) and *Fanca*^-/-^ mice. At the molecular level, we found that FLASH reduced unresolved DNA damage in basal epithelial cells post-irradiation. Together these findings highlight the potential for FLASH to reduce radiation-induced oral mucositis in the FA genomic background.

## Materials and Methods

### Study Design

The objective of this study was to investigate the normal tissue-sparing effect of FLASH compared to conventional dose rate (CONV) irradiation to the oral cavity in *Fanca*^*-/-*^ mice. All procedures for animal care and use were approved by the Institutional Animal Care and Use Committee of Stanford University in accordance with institutional and NIH guidelines. Cohorts of *Fanca*^+/+^ and *Fanca*^-/-^ mice (129/Sv) [27] were generated from *Fanca*^+/+^ and *Fanca*^-/-^ breeding pairs [20] and irradiated with both FLASH and CONV modalities. Male mice aged 10-12 weeks were anesthetized with a mixture of ketamine (100 mg/kg) and xylazine (10 mg/kg) by intraperitoneal injection. Control mice were similarly anesthetized but did not receive radiation. The toxicity of FLASH compared to CONV was evaluated in non-tumor bearing mice. *Fanca*^+/+^ and *Fanca*^-/-^ mice were irradiated with 26 and 18 Gy, respectively, of FLASH or CONV, and tongues were harvested at 12 hours (hpi) and 10 days (dpi) post-irradiation. Radiation dose and tissue-harvesting timepoint were selected based on pilot study data, with genotype-specific doses used to optimize visualization of radiation-induced ulceration while minimizing mortality in *Fanca*^+/+^ and the more radiosensitive *Fanca*^-/-^ backgrounds (Supp. Fig. 1). Tongues were stained with toluidine blue and sectioned for histological analysis of oral mucositis. Immunohistochemistry was also performed on tongue sections to evaluate toxicity on a molecular level.

*Additional methods are located in the Supplemental Materials.*

## Results

### FLASH Irradiation Decreases Radiation-Induced Oral Mucositis in *Fanca*^+/+^ and *Fanca*^-/-^ Mice

To compare the toxicity of FLASH and conventional oral cavity irradiation in the *Fanca*^+/+^ and *Fanca*^-/-^ backgrounds, we used a clinical linear accelerator to deliver reproducible and homogenous lateral and depth dose throughout the mouse oral cavity in FLASH (190 Gy/s) and CONV (0.2 Gy/s) modalities (Fig. 1A-E).

**Figure 1.**
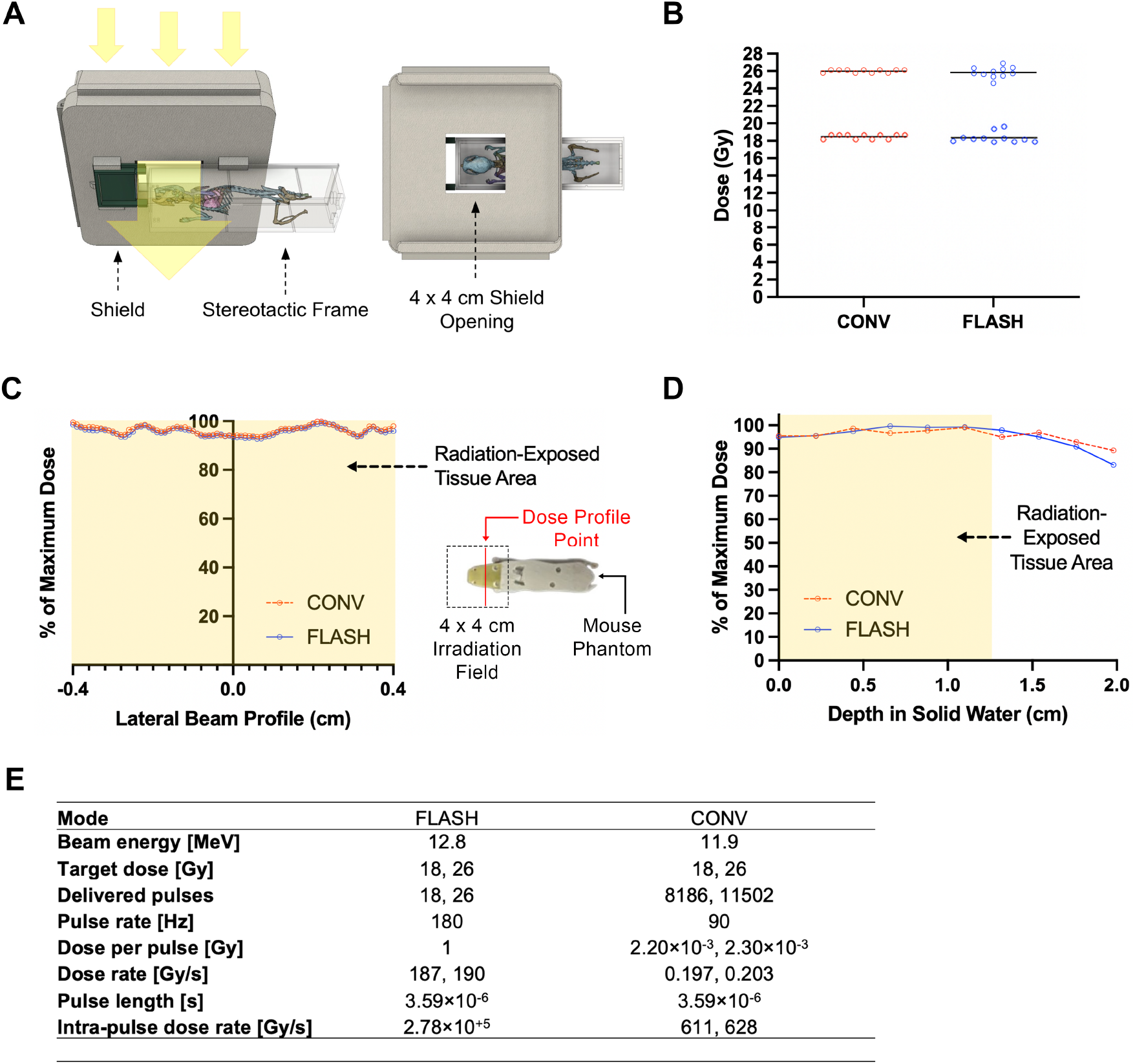
Experimental configuration for ultra-high dose rate FLASH and conventional oral cavity irradiation. (A) 3-D design of irradiation shield and stereotactic frame for reproducible mouse positioning within the irradiation field. Shield consists of 3-D printed PLA plastic shell filled with 1.5 cm thick Cerrobend to limit dose delivery from a nominal 12 MeV electron beam to within shield opening. 4 cm × 4 cm shield opening encompasses the entire oral cavity region, extending from the nose to the first rib. (B) Measured doses delivered, which match target values of 18 and 26 Gy for both FLASH and CONV. (C-D) Lateral and dorsoventral beam profiles recorded using radiochromic films. Highlighted regions indicate the projected tissue area within the irradiation field. Doses are uniformly distributed within the oral cavity region. (C) Dose readings within the irradiation field (dashed box) were acquired at indicated positions (red line) on radiochromic film placed in a coronally-sectioned 3-D printed mouse phantom during irradiation. (E) Geometric and beam parameters for FLASH and CONV. Higher pulse rate and dose per pulse in FLASH results in an average dose rate ∼940 times that of CONV.

To evaluate the radiosensitivity of *Fanca*^-/-^ mice compared to *Fanca*^+/+^ controls, we first performed a pilot study using oral cavity irradiation. Our analysis focused on radiation-induced oral mucositis (RIOM), as it represents a primary dose-limiting toxicity in HNSCC patients characterized by acute pain, dysphagia, opioid dependence, and weight loss [34]. Preliminary toxicity assays demonstrated that *Fanca*^-/-^ mice exhibit increased sensitivity to oral irradiation; specifically, 75% (6/8) of the *Fanca*^-/-^ mice met early euthanasia criteria within 10 days post 26 Gy irradiation, whereas 0% (0/8) of the *Fanca*^+/+^ group reached those endpoints (Supp. Fig. 1). Furthermore, we found that 18 Gy in *Fanca*^-/-^ mice induced a severity of oral mucositis comparable to 26 Gy in *Fanca*^+/+^ mice, without resulting in acute morbidity (Supp. Fig. 1). Therefore, we utilized 26 Gy for the *Fanca*^+/+^ background and 18 Gy for the *Fanca*^-/-^ background to evaluate and compare the normal tissue effects of FLASH versus conventional oral cavity irradiation.

To investigate the safety of FLASH irradiation to the head and neck region, we delivered 26 or 18 Gy FLASH and CONV to the oral cavity of *Fanca*^+/+^ and *Fanca*^-/-^ mice and analyzed the tongues for toxicity at 10 days post-irradiation (dpi). In the *Fanca*^+/+^ background, FLASH and CONV-irradiated mice demonstrated a similar decline in body weight following irradiation (Fig. 2A-B). Notably, in the *Fanca*^-/-^ background, 4/7 FLASH-irradiated mice started to gain weight by 10 dpi, while all CONV-irradiated mice continued to lose weight (Fig. 2A-B). Oral ulceration severity, quantified as the fraction of toluidine-blue stained tongue area relative to total tongue area, was reduced with FLASH compared to CONV in both *Fanca*^+/+^ and *Fanca*^-/-^ mice (Fig. 2C-D). Histological analyses of the tongues using an established scoring system revealed *Fanca*^+/+^ CONV-irradiated tongues scored higher for mucositis severity with 100% of mice with a grade 3 or 4 lesion compared to 43% of mice that received FLASH irradiation (Fig. 2E-F, Sup. Fig. 3). In the *Fanca*^-/-^ mice, 100% of CONV-irradiated tongues developed a grade 3 or 4 lesion compared to 57% of FLASH-irradiated tongues that developed a grade 3 or 4 lesion. Together these data indicate that FLASH irradiation reduces radiation-induced tongue injury in both *Fanca*^+/+^ and *Fanca*^-/-^ mice compared to CONV irradiation.

**Figure 2.**
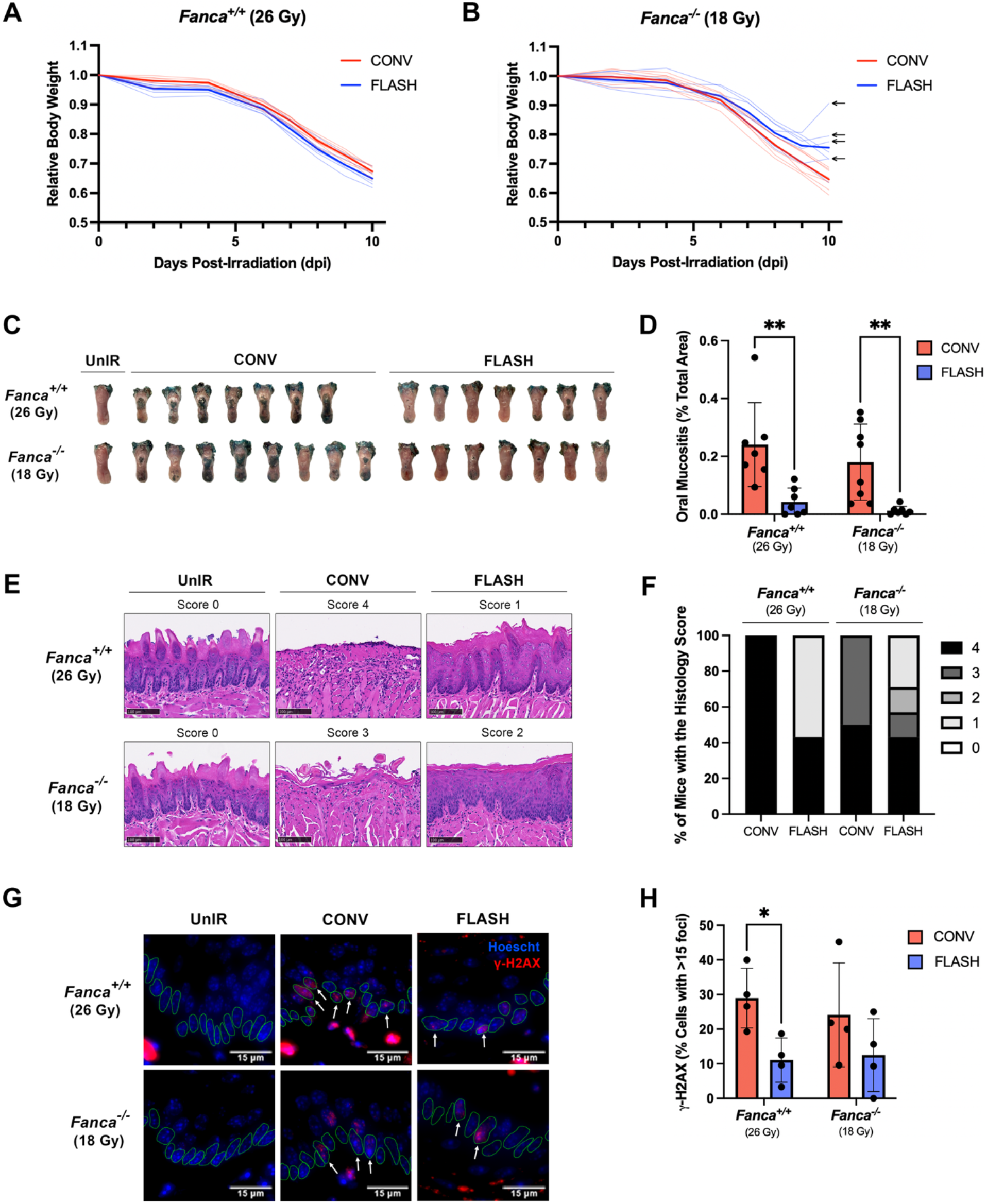
FLASH reduces oral mucositis compared to CONV in *Fanca*^+/+^ and *Fanca*^-/-^ mice. *Fanca*^*+/+*^ and *Fanca*^-*/-*^ mice were irradiated with 26 and 18 Gy, respectively, of CONV or FLASH and tongues were harvested at 10 dpi. For *Fanca*^+/+^, UnIR n=5, CONV n=7, and FLASH n=7. For *Fanca*^-/-^, UnIR n=5, CONV n=8, and FLASH n=7. (A-B) Relative body weight over time after irradiation at 0 dpi; bold lines represent the averages for each cohort. Arrows indicate mice that gained weight at 10 dpi. (C) Tongues were stained with toluidine blue to visualize ulceration. Not all UnIR tongues are pictured. UnIR = Unirradiated. dpi = days post-irradiation. (D) Ulcerated area expressed as a percentage of total tongue area, quantified in digital photographs using Fiji ImageJ. Each data point represents the average of measurements from three independent scorers (** p < 0.01). UnIR tongues exhibited no ulceration. (E) Representative tongue epithelium images from H&E stained tongue sections. Higher scores indicate more severe toxicity. (F) Percentage of mice in each group with a maximum histopathologic score of 0, 1, 2, 3, or 4 (** p < 0.01). (G) Representative γ-H2AX (red) and Hoesch (blue) stained images of tongue epithelia from irradiated and unirradiated *Fanca*^+/+^ and *Fanca*^-/-^ mice at 12 hours post-irradiation (hpi). Arrows indicate basal epithelial cells (outlined in green) with >15 γ-H2AX foci. (H) Quantification of positive γ-H2AX staining, represented by the percentage of total basal epithelial cells with >15 punctate γ-H2AX foci. n=4 mice per group. Unirradiated tongues exhibited no positive γ-H2AX basal epithelial cell staining. An average of 31 basal epithelial cells were analyzed per section. (* p < 0.05).

### FLASH Reduces Unresolved Radiation-Induced DNA Damage in the Tongue Epithelium

The proliferative basal cell layer of the tongue epithelium is essential for epithelial maintenance and regeneration following injury and is particularly radiosensitive due to its high mitotic activity [35]. Consequently, damage in this compartment disrupts tissue repair, contributing to acute and chronic oral toxicities. Therefore, we investigated whether FLASH reduces basal cell DNA damage relative to CONV as a potential mechanism underlying its reported normal tissue-sparing effects.

At 12 hours post-irradiation (hpi), regardless of genetic background, unresolved DNA damage was evident in tongue basal epithelial cells of both CONV and FLASH-irradiated mice, as indicated by γ-H2AX staining, whereas unirradiated controls showed no detectable signal (Fig. 2G). Within *Fanca*^+/+^ mice, the percentage of basal epithelial cells containing greater than 15 punctate γ-H2AX foci was increased in CONV-irradiated tongues compared to FLASH-irradiated counterparts (Fig. 2H). *Fanca*^*-/-*^ mice demonstrated a similar trend, with CONV-irradiated tongues exhibiting a higher percentage of γ-H2AX positive basal epithelial cells compared to FLASH-irradiated counterparts, although the difference did not reach statistical significance. These data suggest that FLASH irradiation reduces unresolved DNA damage burden compared to CONV irradiation in the basal cell layer of the tongue epithelium.

## Discussion

Here we demonstrate that FLASH mitigates radiation-induced oral mucosal toxicity in both *Fanca*^+/+^ and *Fanca*^-/-^ mice, indicating that the normal tissue-sparing properties of FLASH may extend to the setting of intrinsic radiosensitivity.

Oral mucositis in the tongue epithelium is driven largely by injury to the proliferative basal epithelial compartment, where radiation-induced cellular injury, including DNA damage, disrupts epithelial renewal and impairs barrier integrity. In line with this pathology, we observed reduced ulceration, improved histopathologic architecture, and lower mucositis severity scores in *Fanca*^+/+^ mice and *Fanca*^-/-^ mice following irradiation with FLASH compared to CONV. Consistent with our findings in *Fanca*^+/+^ mice using a high-energy electron beam, a recent preclinical study using a proton beam demonstrated reduced tongue ulceration and improved histopathologic architecture with FLASH compared to standard dose rate proton therapy [8], suggesting that this effect is preserved across radiation types. Importantly, the phenotypic differences observed in the present study were accompanied by a reduction in γ-H2AX burden within basal epithelial cells at 12 hours post-irradiation (hpi) with FLASH. As γ-H2AX serves as a surrogate marker of DNA double-strand breaks, the attenuation of unresolved γ-H2AX signal suggests that a reduction of initial damage induction and/or enhanced DNA repair processes may contribute to the observed tissue sparing with FLASH. These findings align with prior reports demonstrating reduced unresolved γ-H2AX in normal tissues following isodoses of FLASH relative to CONV [2,36], extending this observation to the basal epithelial compartment of the tongue. In contrast, reports examining earlier timepoints have yielded mixed results, with some studies observing reduced γ-H2AX signaling following FLASH while others report no significant differences relative to CONV [36,37].

Notably, FLASH reduced oral mucositis in the *Fanca*^-/-^ background. *FANCA* plays a central role in the Fanconi anemia DNA repair pathway, and its loss confers heightened radiosensitivity due to impaired repair of DNA lesions [12]. FA proteins collaborate in the nucleus to coordinate multiple repair processes including DNA lesion recognition, incision, bypass, and repair [12,24], and have been shown to interact with other DNA repair pathways such as nucleotide excision repair, translesion synthesis, and alternative end joining [25,26]. Our findings demonstrate that FLASH irradiation maintains its ability to reduce the severity of radiation-induced oral mucositis in this background, indicating that the FLASH effect may not rely exclusively on FA-mediated repair processes and may therefore retain tissue-sparing potential even in intrinsically radiosensitive settings. The lack of statistical significance in FLASH-mediated γ-H2AX reduction within the *Fanca*^-/-^ cohort may stem from constraints in dose and timing rather than an absence of biological effect. While doses were empirically selected based on pilot data to induce comparable levels of tongue ulceration across genotypes, these parameters may not be optimal for resolving differences between FLASH and CONV for secondary endpoints. Future studies are needed to determine whether alternative dosing or a more granular temporal analysis of DNA repair kinetics in the *Fanca*^-/-^ background would reveal significant divergence. Furthermore, it remains to be determined whether FLASH-mediated radioprotection in the oral mucosa is driven by mechanisms independent of traditional DNA damage and repair pathways.

Collectively, these findings underscore the translational significance of FLASH in intrinsically radiosensitive contexts. Patients with inherited DNA repair deficiencies, including those with Fanconi anemia, experience profound toxicity following conventional radiotherapy and often face limited therapeutic options. Strategies that expand the normal therapeutic window without compromising tumor control are therefore of particular importance in this population. Future studies incorporating tumor-bearing models will be critical to determine whether the normal tissue-sparing effects observed here are maintained without loss of anti-tumor efficacy. Taken together, our data support continued preclinical evaluation of FLASH as a strategy to mitigate radiation-induced normal tissue injury in radiosensitive settings.

## Conclusion

FLASH irradiation reduces radiation-induced oral mucosal toxicity in *Fanca*^+/+^ and *Fanca*^-/-^ mice. This tissue-sparing effect is accompanied by reduced unresolved DNA damage in the basal epithelial compartment of the tongue, a critical site for epithelial regeneration. Together, these findings support the translational potential of FLASH as a strategy to preserve oral mucosa in patients, including intrinsically radiosensitive populations, where conventional radiotherapy is often dose-limiting.

## Acknowledgements

This work was supported by NCI grants P01CA244091 (BWL).

## Author Contributions

Conceptualization PL, ME, JG, BWLJ, EBR

Data Curation PL, EBR

Formal analysis PL, MP, MZ, SM, DC, LW, SR, KMC, EEG

Funding acquisition BWLJ, EBR

Investigation PL, MP, MZ, SM, DC, LW, SR, KMC, EEG

Methodology PL, MP, MZ, SM, DC, LW, SR, KMC, EEG

Project administration ME, JG, BWLJ, EBR

Resources ME, JG, BWLJ, EBR

Supervision FD, EEG, ME, JG, BWLJ, EBR

Validation BWLJ, EBR

Visualization PL, EBR

Writing-original draft PL, EBR

Writing-review and editing PL, MP, MZ, SM, DC, LW, SR, KMC, EEG, SR, FD, EEG, ME, JG, BWLJ, EBR

