## Supplemental Methods and Figures for "FLASH reduces radiation-induced oral mucositis in a mouse model of Fanconi anemia"

### **Supplemental Materials and Methods**

*DNA Isolation and Mouse Genotyping*

*Fanca^-/-^* mice and littermate controls were bred in house as described elsewhere [28], and were generously provided by Dr. Markus Grompe (Oregon Health & Science University, Portland, OR). DNA was isolated for genomic PCR from mouse tails according to the protocol of Laird et al. [29]. Wild-type and mutant *Fanca* were detected using the primers in Table 1 (Supp. Fig. 2). Genotyping analysis of all mice in the study confirmed that mice were either *Fanca*^+/+^ or *Fanca*^-/-^ (Supp. Fig. 2).

| Forward Primer | 5’-TTCCTTCAAAGCTGCTGGGG-3’ |
| --- | --- |
| Reverse WT Primer | 5’-CAGTGACATCTTCCTTCCTAACTCC-3’ |
| Reverse Mutant Primer | 5’-GGTGAACGTTACAGAAAAGCAGGCT-3’ |

**Table 1.** Wild-type and mutant *Fanca* primer sequences

*Animal Care*

Mice were housed in standard-sized (M-BTM) Innovive mouse cages on ventilated Innovive cage racks, with 1-5 mice per cage. Each cage included 1 cm depth of ALPHA-dri bedding (Shepherd Specialty Papers, Watertown, TN) and approximately 6-8 g of Enviro-dri nesting material (Shepherd Specialty Papers, Watertown, TN). Mice were provided ad libitum with a 2018 Teklad 18% protein rodent diet (Envigo, Madison, WI) and water was provided via Aquavive Mouse Chlorinated Water Bottles (Innovive, San Diego, CA).

*Mouse Irradiation*

We developed an irradiation shield consisting of a 3-D printed PLA plastic shell filled with a 1.5 cm thick layer of Cerrobend to shape the electron beam and minimize dose leakage from bremsstrahlung radiation generated in the shield materials. The shield contains a central opening measuring 4 cm by 4 cm to define the irradiation field. Mice were positioned in a custom mouse stereotactic positioning frame 3-D printed from PLA plastic, as previously described [2], with reproducibility ensured by fixing the front teeth on a metal wire. The stereotactic frame slides into the shield such that the irradiation field extends from the tip of the nose to the first rib, encompassing all of the oral cavity. Calibration using radiochromic film (Ashland Advanced Materials, Bridgewater NJ) dosimetry performed in solid water and an anatomically realistic 3-D printed mouse phantom [30] was used to correlate delivered dose with AC current transformer (ACCT) charge measurements [31], and to confirm accurate dose delivery within the desired 4 cm by 4 cm field of irradiation. This calibration enabled dose verification during subsequent irradiation experiments using ACCT readouts.

We used a Varian TrueBeam radiotherapy system (Varian Medical Systems, Palo Alto, CA) as previously described [31] to deliver both FLASH and CONV irradiation with a nominal 12 MeV electron beam. FLASH was delivered at a dose per pulse of 1 Gy and a pulse repetition rate of 180 Hz for an average dose rate of approximately 190 Gy/s (for 18 or 26 pulses). CONV was delivered at a dose per pulse of approximately 2E-3 Gy and a pulse rate of 90 Hz for an average dose rate of approximately 0.2 Gy/s. Delivery of the target doses of 18 and 26 Gy for both FLASH and CONV was confirmed by ACCT readouts. A single irradiation session was performed for each cohort: tissues collected 10 days post-irradiation were irradiated between approximately 16:00 and 19:00, and tissues collected 12 hours post-irradiation were irradiated between approximately 20:00 and 22:00. Mice were anesthetized 5-10 minutes prior to irradiation. During irradiation, they were maintained on room air without external warming. After irradiation, mice were returned to their cages, which were positioned partially on heating pads to facilitate recovery from anesthesia until they regained consciousness. Mice were then housed in their standard housing environment and were monitored daily for early euthanasia criteria that include body weight, appearance, respiratory rate, general behavior, and provoked behavior according to the Mouse Intervention Scoring System (MISS 3; [32] ).

*Tissue Processing and Histological Analysis*

Animals were euthanized by CO_2_ asphyxiation. Tongues were harvested 12 hours and 10 days post-irradiation and immersion-fixed in 10% neutral buffered formalin for 24 hrs, then PBS for 24 hrs, and stored in 70% ethanol. Six transverse sections were collected across the length of each tongue. Formalin-fixed tissues were processed, embedded in paraffin, sectioned at 5 μm, and stained with hematoxylin and eosin.

*Evaluation of Oral Mucositis*

Tongues that were harvested at 10 days post-irradiation were submerged in a solution of 1% toluidine blue in 10% acetic acid. Tongues were then rinsed with 10% acetic acid until no further recovery of dye. Area of ulceration was quantified as a percentage of total tongue area with deep blue staining using the Fiji ImageJ software to analyze digital photographs and averaged across three independent scorers. Tongues that were sectioned and processed for histology were scored for oral mucositis according to the histopathologic criteria published by Sunavala-Dossabhoy et. al [33] (Supp. Fig. 3). In addition, individual mouse body weights were monitored every other day from 0-6 days post-irradiation (dpi), followed by daily measurements from 7-10 dpi.

*γ-H2AX Immunofluorescence*

Slides with paraffin-embedded tongue sections were baked, deparaffinized in xylene and graded alcohol series, and subjected to antigen retrieval in 10 mM sodium citrate buffer. Tissue sections were blocked with the Mouse On Mouse (M.O.M.) blocking reagent for 30 min (Vector Labs) and serum-free protein block (Dako) and subsequently incubated with mouse mAb for Phospho-Histone H2A.X (Ser139, 9F3, Abcam; 1:1000) at 4 °C in a humidified chamber overnight. Slides were washed using 1X PBS and incubated with Alexa Fluor 594 conjugated anti-mouse antibody (Invitrogen Cat#A32742; 1:1000) for 30 min at 37 °C. Antibodies were diluted using 1X PBS containing 0.1% BSA and 0.2% Triton X-100. Slides were then counterstained with Hoechst (1.2 ug/mL; Thermo Scientific Cat#62249). Z-stack images were acquired using a Leica DMi8 fluorescence microscope, capturing 31 slices at a step size of 0.4 μm (z-direction) to encompass the entire nucleus as previously described [2]. One 60X image of the dorsal region of the tongue adjacent to the midline was collected per mouse which correlates to the region of maximum ulceration observed with histopathologic analysis. γ-H2AX foci were evaluated in maximum intensity projections generated from the acquired Z-stacks. An average of 31 basal epithelial cells were analyzed per section, and punctate γ-H2AX foci were quantified on a per-cell basis using Fiji ImageJ software (n = 4-5 mice/group). For each mouse, we scored the proportion of cells containing 15 or more foci.

*Statistical Analysis*

Due to a difference in radiation dosing between genotypes, statistical comparisons were restricted to FLASH versus CONV treatment within each genotype and group comparisons were performed using a two-tailed Mann–Whitney test. Body weights were analyzed using a two-way repeated-measures ANOVA. Post hoc analysis included pair-wise comparisons using a Tukey adjustment for multiple comparisons. All analyses were conducted in Prism v 10.2.0 (GraphPad Software, San Diego, CA).

### **Supplementary Figures**

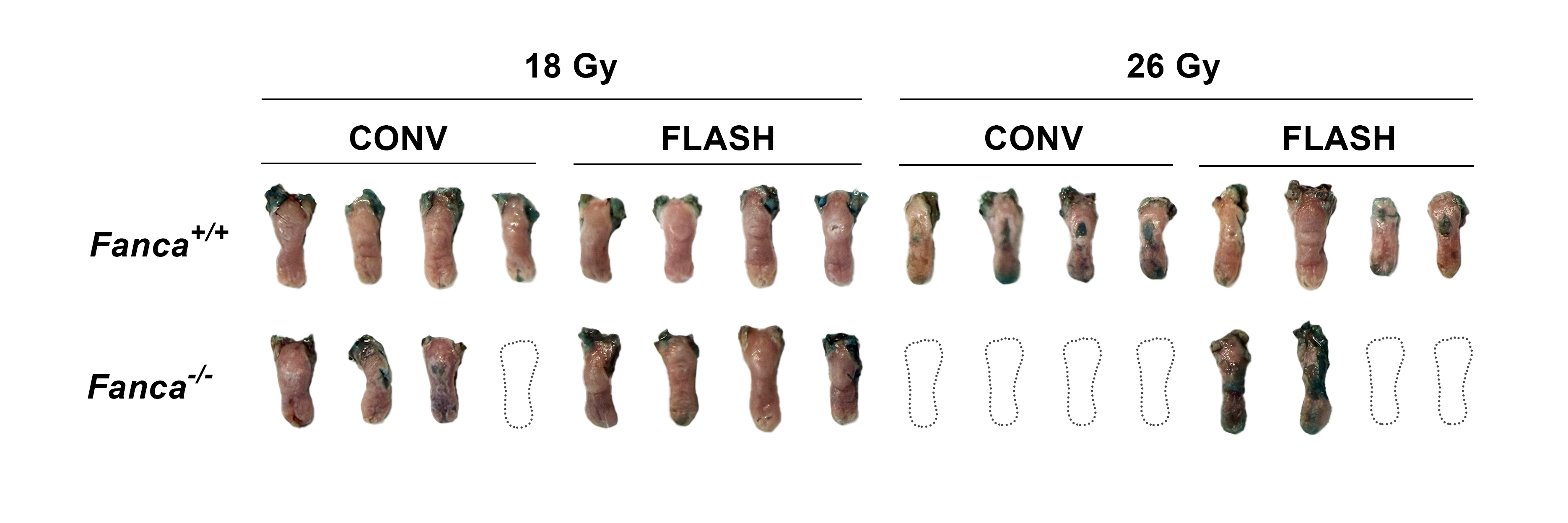

**Supplementary Figure 1. Pilot evaluation of oral mucositis in FLASH and CONV-irradiated tongues.** *Fanca*^+/+^ and *Fanca*^-/-^ tongues were irradiated with either 18 or 26 Gy using FLASH or CONV and harvested at 10 dpi, then stained with toluidine blue to visualize ulceration (n=4 mice per group). Mice that died prior to the 10-dpi harvest timepoint are indicated by dotted outlines. *Fanca*^-/-^ mice exhibited high mortality at 26 Gy, whereas *Fanca*^+/+^ mice showed minimal ulceration at 18 Gy. Consequently, different doses were selected for each genotype, with *Fanca*^+/+^ mice to receive 26 Gy and *Fanca*^-/-^ mice to receive 18 Gy to optimize visualization of radiation-induced ulceration.

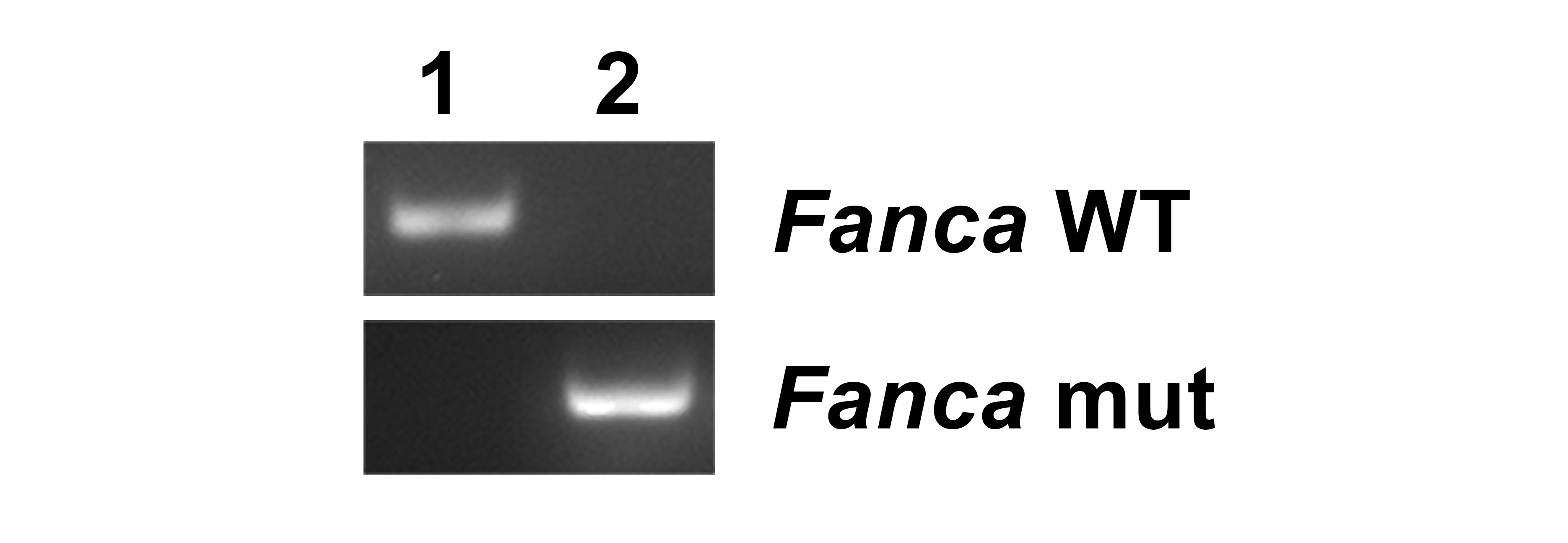

**Supplementary Figure 2. Genotyping of *Fanca*^+/+^ and *Fanca*^-/-^ mice.** Genomic DNA isolated from tails of (1) *Fanca*^+/+^ and (2) *Fanca*^-/-^ mice. Mouse genotypes were confirmed using PCR analysis.

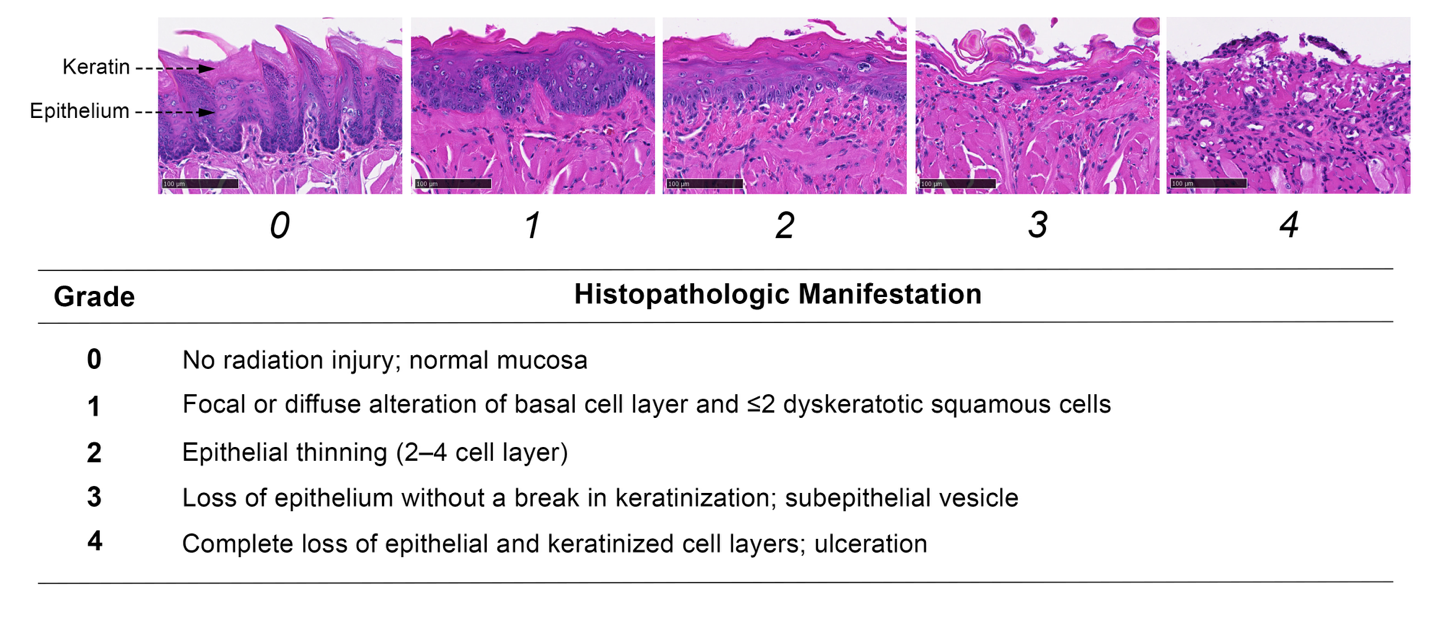

**Supplementary Figure 3. Histopathologic criteria for evaluation of oral mucositis.** Representative histological tongue epithelium images for each histopathologic score, accompanied by scoring criteria adapted from Sunavala-Dossabhoy et. al [33]. Epithelium (comprising stratum basale, stratum spinosum, and stratum granulosum) and keratin (stratum corneum) are indicated.
